# Analytical solutions of the chemical master equation with bursty production and isomerization reactions

**DOI:** 10.1101/2021.03.24.436847

**Authors:** Gennady Gorin, Lior Pachter

**Affiliations:** Division of Chemistry and Chemical Engineering, California Institute of Technology, Pasadena, CA, 91125; Division of Biology and Biological Engineering & Department of Computing and Mathematical Sciences, California Institute of Technology, Pasadena, CA, 91125

## Abstract

Splicing cascades that alter gene products post-transcriptionally also affect expression dynamics. We study a class of processes and associated distributions that emerge from a bursty promoter model coupled to a path graph of downstream mRNA splicing, and more generally examine the behavior of finite-activity jump drivers coupled to a directed acyclic graph of splicing with one or more roots. These solutions provide full time-dependent joint distributions for an arbitrary number of species, offering qualitative and quantitative insights about how splicing can regulate expression dynamics. Finally, we derive a set of quantitative constraints on the minimum complexity necessary to reproduce gene co-expression patterns using synchronized burst models. We validate these findings by analyzing long-read sequencing data, where we find evidence of expression patterns largely consistent with these constraints.

## 1 Introduction

Recent advances in the analysis of single-cell RNA sequencing [1] and fluorescence microscopy [2] enable the quantification of pre-mRNA molecules alongside mature mRNA. These techniques provide an opportunity to infer the topologies and biophysical parameters governing the processes of mRNA transcription, splicing, export, and degradation in living cells. In particular, they provide a novel approach to inferring and studying the dynamics of splicing cascades [3].

However, drawing mechanistic conclusions from transcriptomics requires overcoming numerous statistical and computational challenges. Living cells contain mRNA in low copy numbers, and transient nascent species are even less abundant, leading to potential pitfalls if the discrete nature of the data is not appropriately modeled [4]. One approach to modeling dynamics from count data is to utilize detailed Markov models based on the chemical master equation (CME) [5–7]. Such modeling can, in principle, yield analytical solutions for the dynamics of genes affected by arbitrary splicing networks, thus circumventing tractability problems arising with matrix- and simulation-based methods that can be impractical for large numbers of species [8]. Several classes of analytical solutions to the CME are available [9], but their derivation tends to be *ad hoc*, with limited generalization to more complex systems.

Starting with the approach of Singh and Bokes to the problem of bursty transcription coupled to nuclear export and degradation of mRNA [10], we develop a class of solutions for gene dynamics affected by arbitrary splicing networks under the physiologically relevant assumption of bursty production of mRNA [11]. Fundamentally, the generating function of a Markov chain describing the evolution of molecules in a splicing cascade can be automatically computed, numerically integrated in time, and then inverted by Fourier transformation at an overall computational complexity of 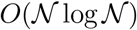 in state space size. We begin with the example of splicing described by a path graph, where the order of intron removal is deterministic, extend the procedure to a much broader class of splicing graphs and driving burst processes, and subsequently demonstrate the existence of the solutions and their isomorphism to a class of moving average processes.

## 2 Methods

### 2.1 Path graph splicing

Consider the system consisting of a bursting gene coupled to a *n*-step birth-death process, characterized by the path graph in Figure 1, where *B* ~ *Geom*(*b*), and all reactions occur after exponentially-distributed waiting times. The bursts occur with rate *k*, the conversion of adjacent transcripts 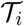 to 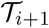 occurs with rate *β*_*i*_, and the degradation of 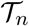 occurs with rate *γ* ≔ *β*_*n*_. We assume the rates of conversion and degradation are all distinct. The amount of species 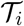 can be described by the non-negative discrete random variable *m*_*i*_. We assume no molecules are present at *t* = 0.

**Figure 1:**
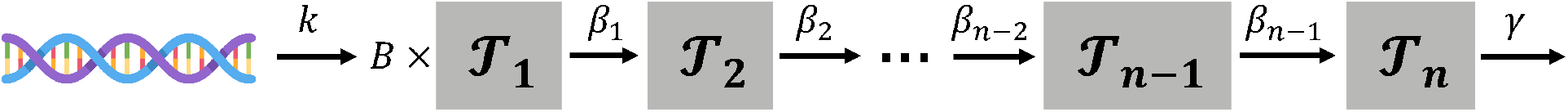
Graph representation of the generic path graph model. The source transcript 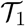 is synthesized at the gene locus in random geometrically distributed bursts (according to a distribution *B* with burst frequency *k*). Each molecule proceeds to isomerize in a chain of splicing reactions governed by successive rates *β*_1_, *β*_2_, ..., *β*_*n*−1_, until reaching the form 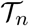, which is ultimately degraded at rate *γ*.

#### 2.1.1 Discrete formulation and algorithm

We would like to compute the probability mass function (PMF) of the count distribution, denoted by *P*(*m*_1_, *..., m*_*n*_, *t*); this corresponds to the probability of observing *m*_1_ molecules of 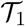, *m*_2_ molecules of 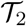, etc., at time *t*. Following a previous derivation [10], this problem can be reframed in terms of a partial differential equation involving the probability generating function (PGF) *G*(*x*_1_, *..., x*_*n*_, *t*):

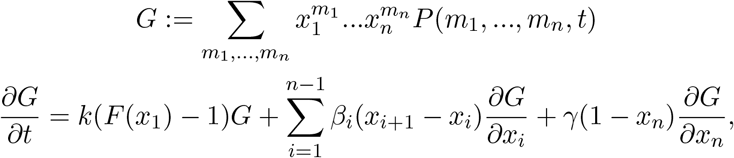

where *F* is the PGF of the burst distribution. Applying the transformations *u*_*i*_ ≔ *x*_*i*_ − 1 and *ϕ* ≔ ln *G* yields the equation:

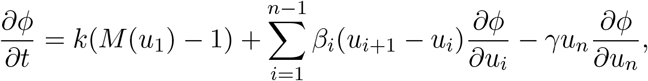

where *M*(*u*) ≔ *F*(1 + *u*). This equation can be solved using the method of characteristics, with formal solution 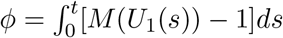. The characteristics *U*_*i*_, *i < n* satisfy 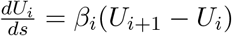, with 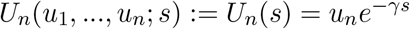.

The functional form of 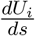 implies that *U*_1_(*s*) is the weighted sum of exponentials 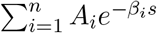 the explicit analytical solution is provided in Section 5. The weights *A*_*i*_ can be computed through a simple iterative procedure, which proceeds from the terminal species and successively incorporates dependence on upstream rates:

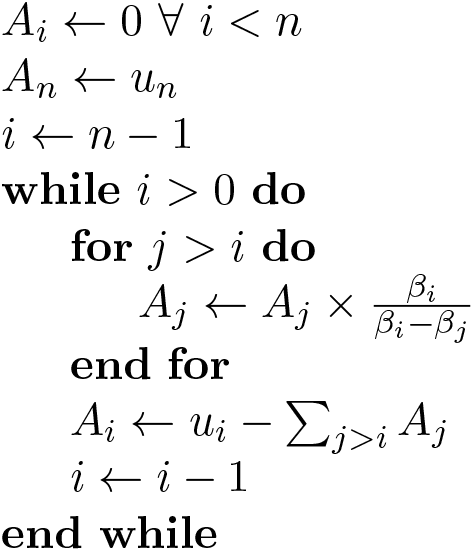

Finally, given the physiologically plausible geometric burst distribution [12], 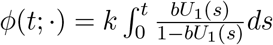. The stationary PGF is *e*^*ϕ*^ evaluated at *t* = ∞; this quantity can be converted to a probability mass function via Fourier inversion. This is an entirely general procedure that can be used for *any* path graph of downstream processing, although regions of stiffness are to be expected, particularly near the degenerate cases of matching rates. In the most general case, any *β*_*i*_ may be equal to any other, leading to a combinatorial explosion of auxiliary degenerate solutions. As the overarching motivation is the statistical inference of biophysical parameters, and the degenerate regions are all of measure zero, there appears to be little physical justification for considering them in further detail. However, we remark that these solution classes are likewise analytically tractable, with mixed polynomial-exponential functional forms entirely analogous to previous work [10].

### 2.2 Directed acyclic graph splicing

The simplicity of the ODEs governing the evolution of the CME lends itself to extensions to the broadest class of graphs representing the stochastic and incremental removal of discrete introns, the directed acyclic graphs.

#### 2.2.1 Example: alternative splicing

Suppose the downstream dynamics are given by a directed rooted tree. The solution procedure is analogous to that used for the path graph. First, starting at the leaves, the path subgraph solutions are produced by the procedure above, yielding a sum of exponentials. Then, at a node of out-degree > 1 (i.e., molecular species with several potential products), the associated ODE has a functional form identical to that of a path graph. Therefore, the solutions are analogous.

As an illustration, consider the simplest tree graph, shown in Figure 2a, where the splicing reactions occur at rates *α*_1_ and *α*_2_ and degradation reactions occur at rates *β*_1_ and *β*_2_. Physically, this graph can be interpreted as a single source mRNA being directly and stochastically converted to one of two terminal isoforms by removal of intron 1 or intron 2. Clearly, 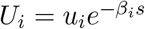 for *i* ∈ {1, 2}.

**Figure 2:**
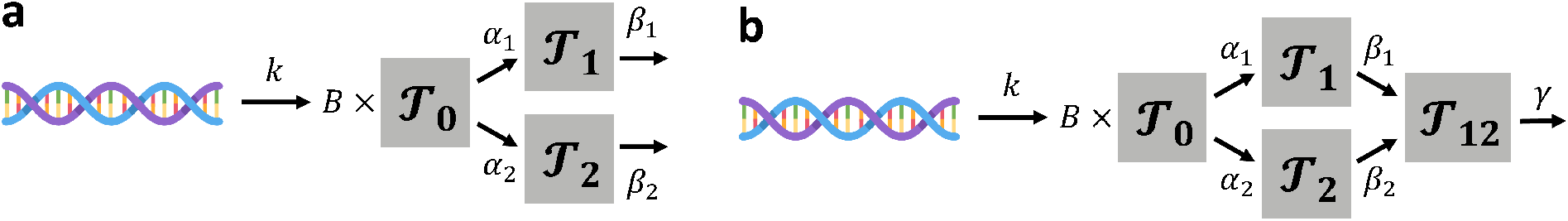
Graph representations of simple directed acyclic graph models. (a) Tree splicing graph with two terminal isoforms. (b) Convergent splicing graph with a single terminal isoform and two intermediate transcripts.

The ODE governing the source species is 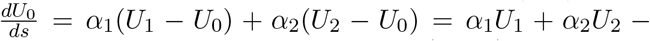 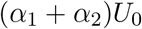. The solution is 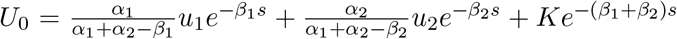, with 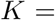 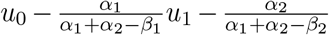. Finally, the expression for *U*_0_ can be directly plugged into the desired burst distribution generating function.

#### 2.2.2 Example: two-intron splicing with non-deterministic order

Consider the same tree graph as in the example above, and suppose 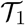 and 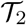 are converted to product 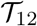 at rates *β*_1_ and *β*_2_, as shown in Figure 2b. Afterward, 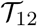 is degraded at rate *γ*. Physically, this graph can be interpreted as a single source mRNA being converted to one of two intermediate isoforms by the removal of one of two introns, then to a single terminal isoform by the removal of the other intron. Clearly, 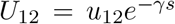. Setting 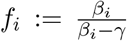, we find 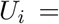 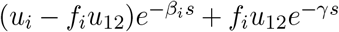. Finally, the dynamics of the source molecule 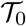 are governed by the following ODE:

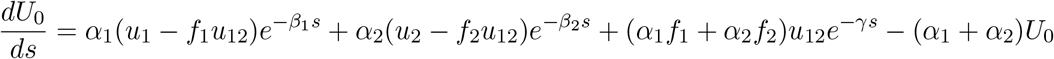

Yet again, the functional form affords a straightforward analytical solution:

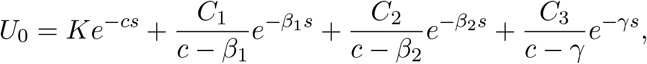

where *c* ≔ *α*_1_ + *α*_2_, *C*_1_ ≔ *α*_1_(*u*_1_ − *f*_1_*u*_12_), *C*_2_ ≔ *α*_2_(*u*_2_ − *f*_2_*u*_12_), and *C*_3_ ≔ *α*_1_*f*_1_ + *α*_2_*f*_2_. From the initial condition *U*_0_(*s* = 0) = *u*_0_, we yield 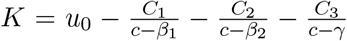. The details of the ODE solution are described in Section 5 and the computation procedure is demonstrated in Figure 3.

#### 2.2.3 Algorithm

This procedure is facile to extend to an arbitrary directed acyclic graph with a unique root. The reaction system is fully characterized by two arrays, the stoichiometric matrix matrix *S* and the rate vector *r* used in the stochastic simulation algorithm [13]. Thus, for example, the path graph can be represented by the full rate vector [*k, β*_1_, *..., β*_*n*−1_, *γ*] and the following stoichiometric matrix:

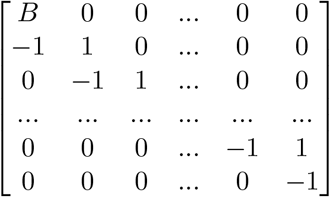

**Figure 3:**
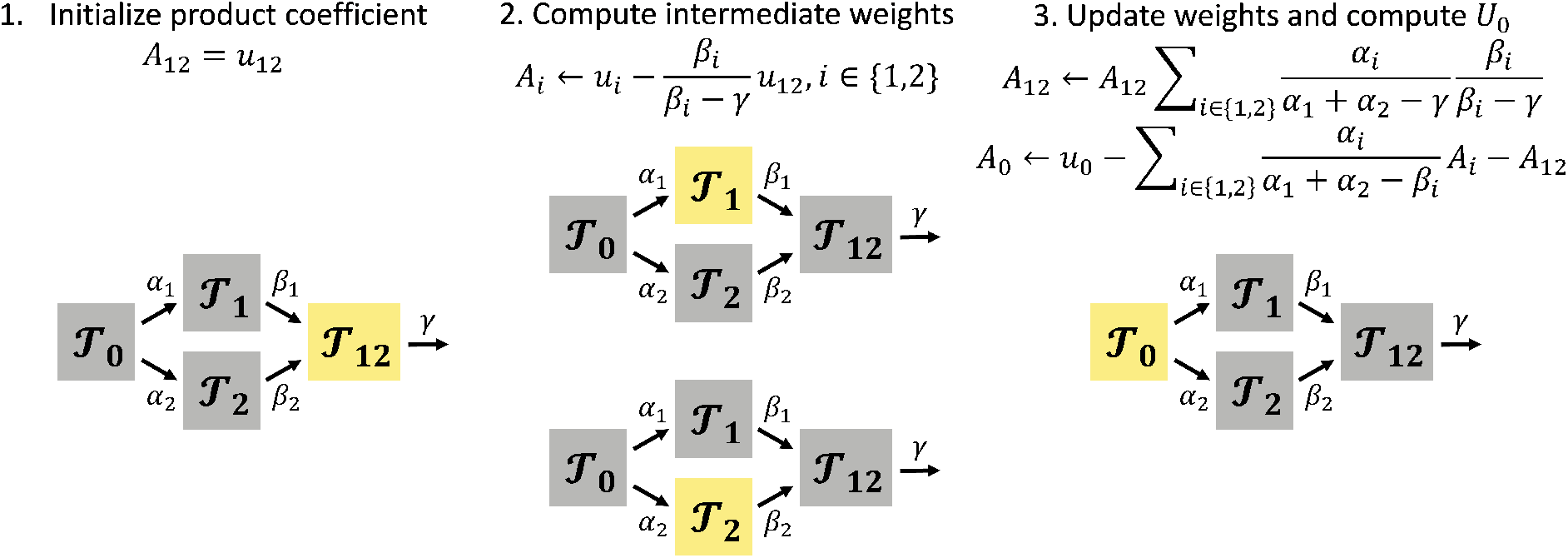
Illustration of the solution algorithm. The differential equation structure requires the backward propagation of downstream species’ solutions, weighted by ratios of rates.

A well-defined system has *n* species and *q* + 1 reactions. We assume unbounded accumulation does not occur and the graph is connected, so each species has influx and efflux reaction pathways, implying *q* ≥ *n*. Further, we suppose that each species is associated with at most one degradation reaction; if several channels are available, they can be added by the superposition property of a Poisson process. Finally, we assume that all isomerization and degradation rates are distinct. Since the underlying graph is acyclic, there exists at least one terminal species with a single efflux reaction. Thus, consider a single production reaction with rate *k* coupled to a set of isomerization reactions with rates *c*_*ji*_ and degradation reactions with rates *c*_*j*0_.

The downstream dynamics are determined by *U*_1_, an exponential sum with *n* terms. In this generic case, *lumped* rate exponents *r*_*i*_ must be computed as the net efflux rate from each molecule. A simple implementation of the routine is outlined below:

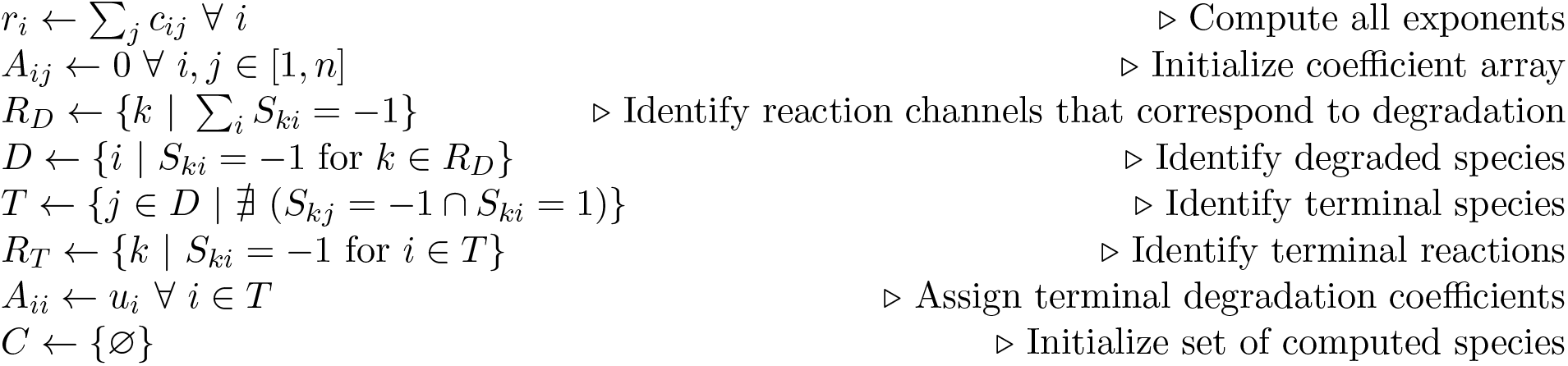
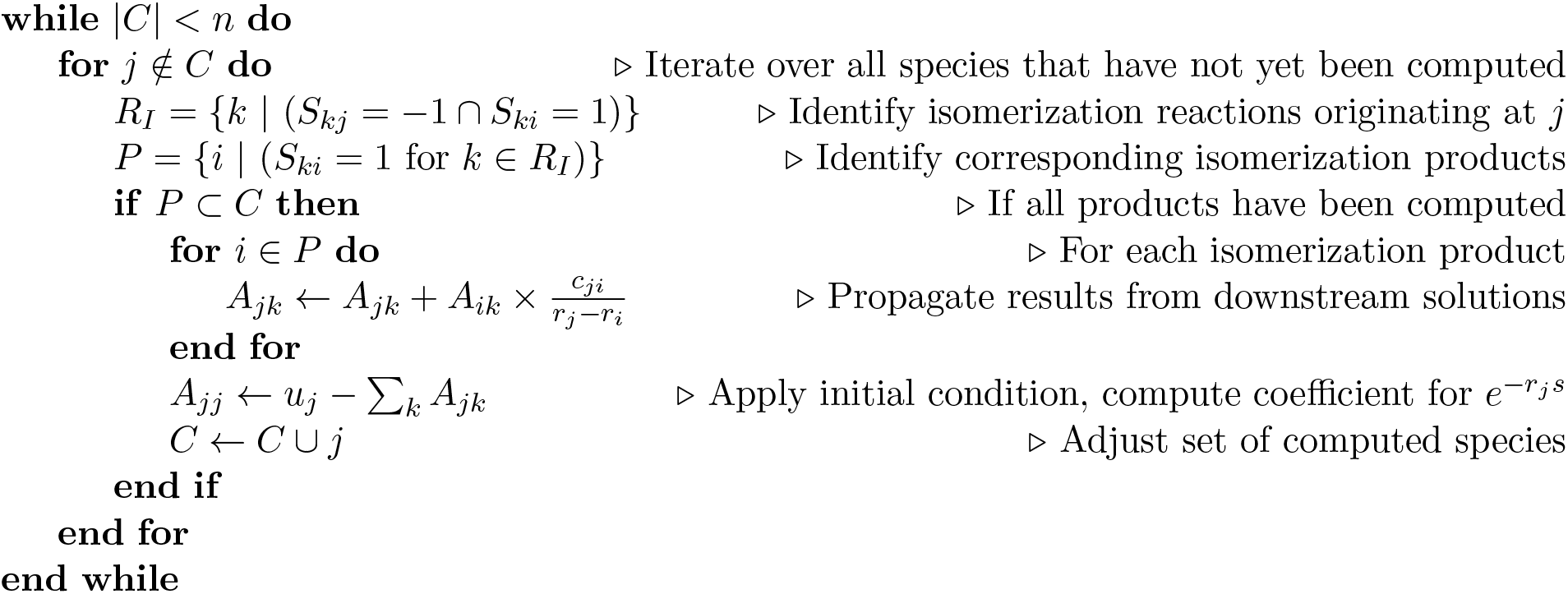

The terminal exponential sum is given by 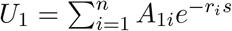. As before, the full time-dependent solution for any burst distribution with a well-defined moment-generating function is computable by quadrature. The case of the geometric burst distribution yet again corresponds to 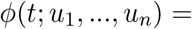 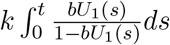.

### 2.3 Continuous formulation

We have defined a series of discrete joint distributions induced by a graph governing splicing and degradation. However, we can equivalently recast the problem in terms of the stochastic differential equations (SDEs) governing the distributions’ Poisson intensities Λ_*i*_. Specifically, the following identity can be used to relate a set of stochastic processes Λ_1_, *...,* Λ_*n*_ with joint distribution 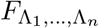 to the solution of the CME [9, 14–16]:

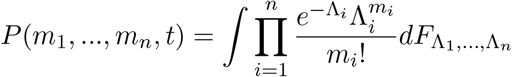

As an illustration, we can consider the intensity of an *n*-step isomerization process driven by a finite-activity compound Poisson Lévy subordinator *L*_*t*_:

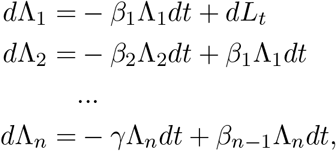

This formulation has an intimate connection with the theory of moving average processes, which can be immediately seen by applying variation of parameters to the Poisson representation:

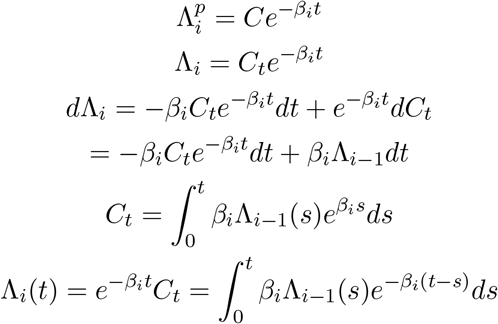

By convention, *β*_*n*_ ≔ *γ* and Λ_0_*ds* ≔ *dL*_*s*_, the underlying driving subordinator.

Thus, each Λ_*i*_ is the exponentially-weighted moving average of Λ_*i*−1_; Λ_1_ is the moving average of the Poisson shot noise introduced by the compound Poisson subordinator. Although ample literature exists on the topic of moving average processes [17], it generally considers the problem of inference and prediction from discrete-time observations. Furthermore, discussions in the context of the chemical systems only tend to consider the first-order moving average of shot noise [9].

This formulation implies that any *n*-step isomerization process is identical to an (*n* − *i*)-step isomerization process driven by an order *i* iterated moving average process. This identity affords a route for the analytical computation of hybrid continuous-discrete system solutions merely by marginalizing the first *i* species.

The Poisson representation enables the simultaneous discussion of the properties of 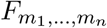, the stationary distribution of the continuous-time Markov chain, and 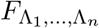, the stationary distribution of the underlying series of stochastic differential equations. In fact, from standard properties of Poisson mixtures [18], the PGF *G*(*x*_1_, *..., x*_*n*_) of the former evaluated at *u*_*i*_ = *x*_*i*_ − 1, precisely yields the moment-generating function (MGF) of the latter.

This connection also allows the evaluation of the solution for a broad class of compound Poisson driving processes. Specifically, 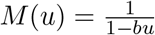 is the MGF associated with exponentially distributed jumps. In the discrete domain, this translates to the geometric burst distribution. However, the downstream dynamics, represented by *U*_1_(*s*), are independent of the burst distribution. Therefore, the procedure affords computable solutions for any finite-activity pure-jump Lévy driving process. Extensions to generic directed acyclic graphs are entirely analogous. These solutions, identical to the CME case, can be used to exactly solve arbitrary DAGs representing *continuous*-valued molecular concentrations. This representation is conventional for high-abundance species [19].

This formulation occasionally enables the confirmation of CME results using standard properties of SDEs. For example, when *L*_*t*_ is a compound Poisson subordinator with exponential jumps, Λ_1_ is the gamma Ornstein–Uhlenbeck process [20–23]. Inspired by a highly general result for constitutive transcription, which states that Poisson distributions always remain Poisson for a birth-death process [14], we may reasonably ask whether equivalent results are available for bursty processes. This intuition turns out incorrect, and straightforward to disconfirm using SDE results. The distribution of the gamma Ornstein–Uhlenbeck process is not gamma for any finite *t* ∈ (0, ∞) – although it does approach a gamma law exponentially fast [21]; therefore, the corresponding Poisson mixture is *not* simply a time-varying negative binomial.

### 2.4 Distribution properties

Using the algorithms above, we can compute the generating functions corresponding to the stochastic processes of interest. However, it is not yet clear that these generating functions are everywhere well-defined. For example, certain physiologically plausible noise models do not have all moments [24]; their generating functions fail to converge in certain regimes. By analyzing the functional form of the downstream process and assuming geometric-distributed bursts, we demonstrate that this class of processes is guaranteed to yield convergent generating functions and finite moments.

#### 2.4.1 Positivity of the exponential sum

First, we demonstrate that the downstream processes yield a strictly positive functional form of time dependence. Noting that the marginal of species *i* yields the functional form 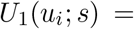 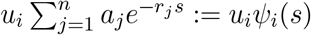, this condition translates to *ψ*_*i*_(*s*) > 0 for all *s* > 0.

Consider *F*(*x*) = *x*, corresponding to constitutive production of the source species (i.e. a Poisson birth process), with no molecules present at *t* = 0. Focusing on the marginal of species *i*, this assumption yields 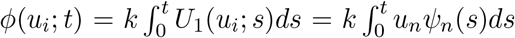. Evaluating *e*^*ϕ*^ at *x*_*i*_ = 0, i.e. *u*_*i*_ = −1, marginalizes over all *j ≠ i* and yields the probability of observing zero counts of species 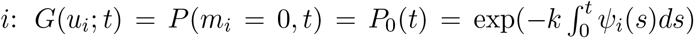. The corresponding time derivative is 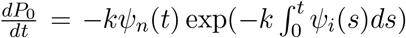. Simultaneously, we know that 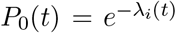 where *λ*_*i*_(*t*) is the solution of the reaction rate equation for species *i* [14]. Clearly, 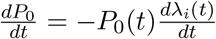. The reaction rate 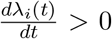 at *t* = 0 under the given initial conditions. Furthermore, 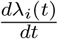 is strictly positive. This follows from the reaction rate equations. By the continuous formulation, *λ*_*i*_ is a weighted moving average of some set of processes {*λ*_*k*_}. *λ*_1_ is a strictly increasing function governed by 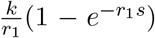. The property of being strictly increasing is retained under moving average and rescaling. Therefore, each successive moving average must be strictly increasing.

Finally, *P*_0_ ∈ (0, 1) *>* 0, because the solution for *m*_*i*_ is given by a Poisson distribution, which has support on all of ℕ_0_. Therefore, 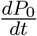 is strictly negative. As the exponential term and *k* are positive, this implies *ψ*_*i*_(*s*) is strictly positive for all *s >* 0.

#### 2.4.2 Existence of generating functions and moments

Next, we show that *G*(*u*_*i*_; *t*), the generating function of the *i*th marginal, is finite for the geometric burst system. This follows from the construction of the original PGF: the marginal PGF is guaranteed to be finite if 1 − *bu*_*i*_*ψ*_*i*_ is never zero. But for the relevant domain 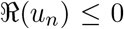, on the shifted complex unit circle, 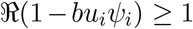, except at the degenerate initial case. The existence of the marginal moments of *m*_*i*_ is implied by the existence of the generating function. The existence of all cross moments follows from the Cauchy-Schwartz inequality. Per standard properties, this existence property holds for both *m*_*i*_ and Λ_*i*_.

The tails of the stationary discrete marginals decay no slower than the geometric distribution. This follows immediately from the lower bound on 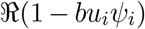, which in turn gives an upper bound on *x*_*i*_ [10]. Equivalently, this follows from the existence of all moments [24]. An analytical radius of convergence has been given previously for *n* = 2 [10], but numerical optimization is necessary to establish rates of tail decay for *n >* 2.

#### 2.4.3 Statistical properties

##### All marginals are infinitely divisible

This follows from the functional form of the PGF: the random variable corresponding to any marginal distribution can be written as as a sum of *q* random variables with burst frequency *k/q*.

##### Only the first marginal is self-decomposable

This follows from the condition that a random variable has a self-decomposable (sd) law if and only if it offers a representation of the form 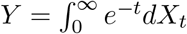, with Lévy *X*_*t*_ [25]. However, only Λ_0_ is Lévy. All downstream intensity processes have nontrivial, almost-everywhere *C*^∞^ trajectories, which implies they cannot be represented by a Lévy triplet: the only permitted continuous Lévy processes are linear combinations of the (non-differentiable) Brownian motion *W*_*t*_ and the trivial process *t*. Therefore, Λ_*i*_ is sd for *i* = 1 and non-sd for all *i >* 1.

##### All stationary marginals are unimodal

Multimodality in the distribution of the moving average over time is contingent on time-inhomogeneity in the trajectory process. However, the underlying driving process is defined to be time-homogeneous, with uniformly distributed jump times; therefore, each downstream process has a unimodal distribution over time. By ergodicity, this distribution is equivalent to the ensemble distribution. Therefore, all marginals of downstream species are unimodal.

#### 2.4.4 Moments

The moments of the marginals can be computed directly from the derivatives of the marginal MGF 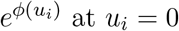

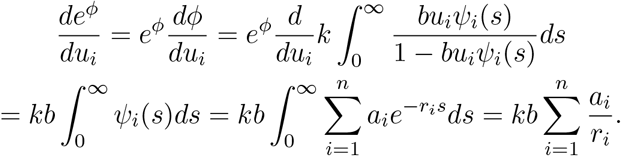

For the path graph system, the CME immediately yields 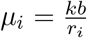. Considering the previously discussed constitutive solution 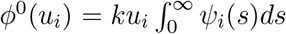, we yield the identity 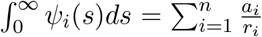, which is equal to 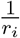 for the path graph system. Per the standard properties of mixed Poisson distributions [26], the value of *μ*_*i*_ is identical for the underlying continuous process and the derived discrete process.

The stationary second moments can be found analogously:

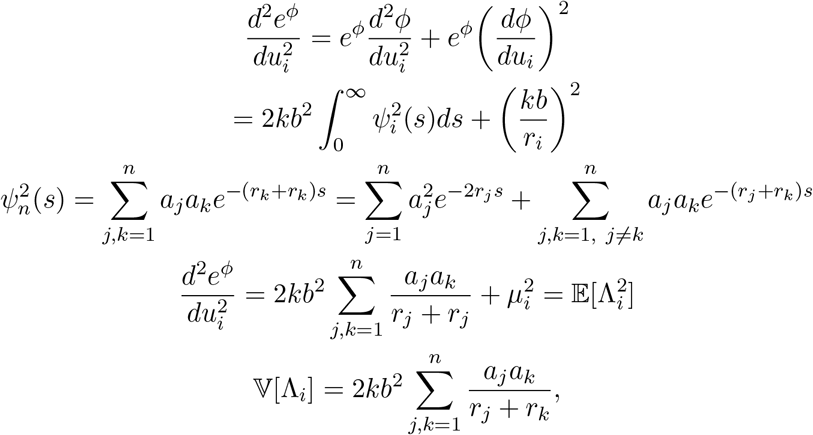

which is straightforward to compute, but does not appear to have an easily amenable analytical form beyond the first few cases. Naturally, the standard properties of Poisson mixtures [26] allow conversion to the discrete domain, with 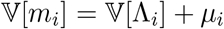.

The covariances can be computed directly from the derivatives of the MGF 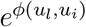, i.e., the marginalized MGF for *u*_*q*_ = 0 for all *q* ≠ *l, i*. The particular functional form of *ϕ*(*u*_*l*_, *u*_*i*_) is given by 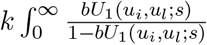. By construction, *U*_1_(*u*_*l*_, *u*_*i*_; *s*) = *u*_*l*_*ψ*_*l*_(*s*)+*u*_*i*_*ψ*_*i*_(*s*), where each *ψ* is the exponential sum corresponding to the marginal of the species in question. This yields:

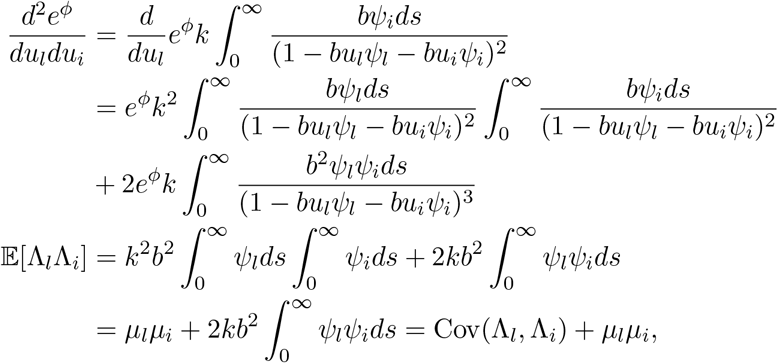

which implies that

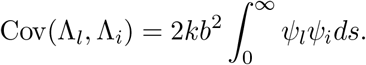

As above,

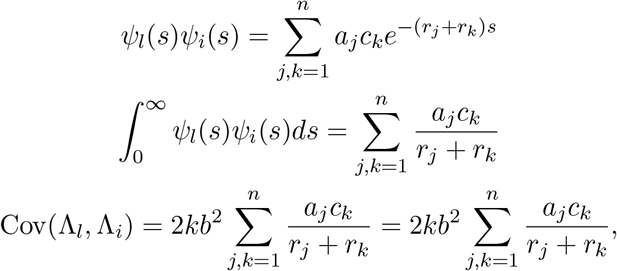

where *a*_*j*_ are the weights associated with *ψ*_*l*_ and *c*_*k*_ are the weights associated with *ψ*_*i*_. This form of the summation is very general; for example, in case of the path graph system with *l > i*, it can be equivalently represented as 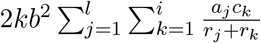. From standard identities, the covariance of a mixed bivariate Poisson distribution with no intrinsic covariance forcing is identical to the covariance of the mixing distribution [26]. Therefore, this result holds for both the CME and the underlying SDE.

The Pearson correlation coefficient follows immediately from the result above:

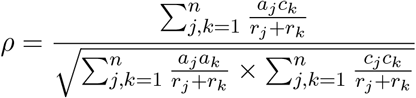

Since mixing decreases variance but not covariance, the correlation coefficient of the discrete system will always be lower than that of its continuous or hybrid analog.

### 2.5 Simulation

To compare the analytical solutions with simulation, we generated a random directed acyclic graph, shown in Figure 4. The numbers of species (7) and isomerization reactions (11) were chosen arbitrarily. We enforced the existence of a single unique source node (a) and the weakly connected property to ensure only a single source mRNA would be present and all isoforms would be reachable from it, but did not impose any other conditions. The number of degraded species (3) was chosen arbitrarily; we assigned degradation reactions to the two sink species (c, e) and randomly chose a degraded intermediate (b) from a uniform distribution over the molecular species.

**Figure 4:**
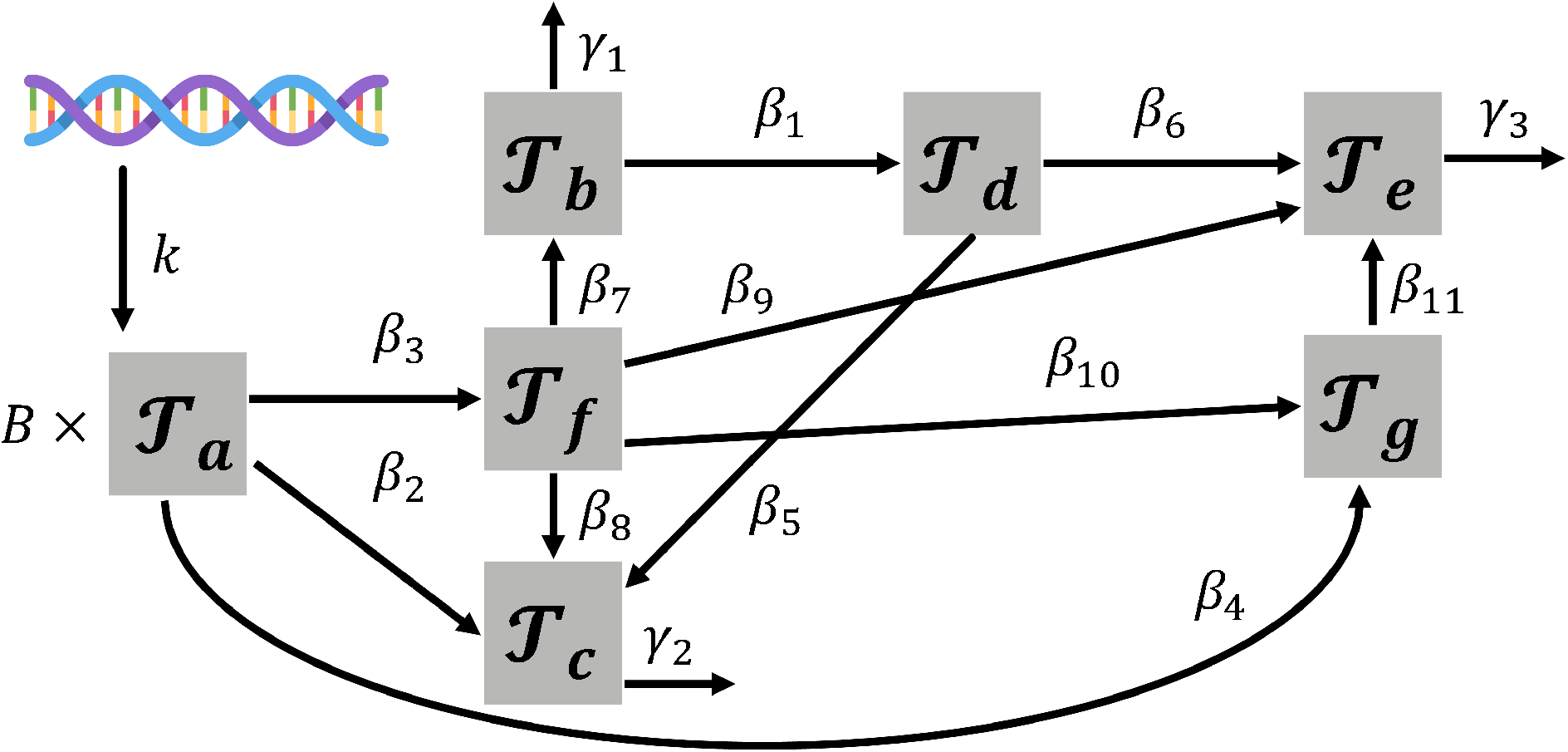
Graph representation of the randomly generated transcription, splicing, and degradation model. A single source isoform 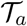 is converted to a variety of downstream isoforms 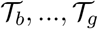, which isomerize according to a directed acyclic graph.

All reaction rates were drawn from a log-uniform distribution on [10^−0.5^, 10^0.5^]; we chose to sample them from a single order of magnitude to avoid the trivial degenerate cases that occur in cases of very slow or very fast export [10]. This process produced the parameter values *k* = 0.44, *β* = [0.48, 2.12, 1.31, 2.21, 1.16, 2.41, 0.4, 1.19, 0.37, 1.19, 0.53], and *γ* = [0.94, 2.38, 0.72], with the indices corresponding to those in Figure 4. Finally, we chose the geometric burst model with *b* = 10.

We applied the algorithm to compute the exponents and coefficients, and computed the stationary distributions of all species. The simulated distributions match the quantitative results for the marginals, as shown in Figure 5. Furthermore, the 49 entries of the covariance matrix are likewise effectively predicted by the procedure for moment calculation.

**Figure 5:**
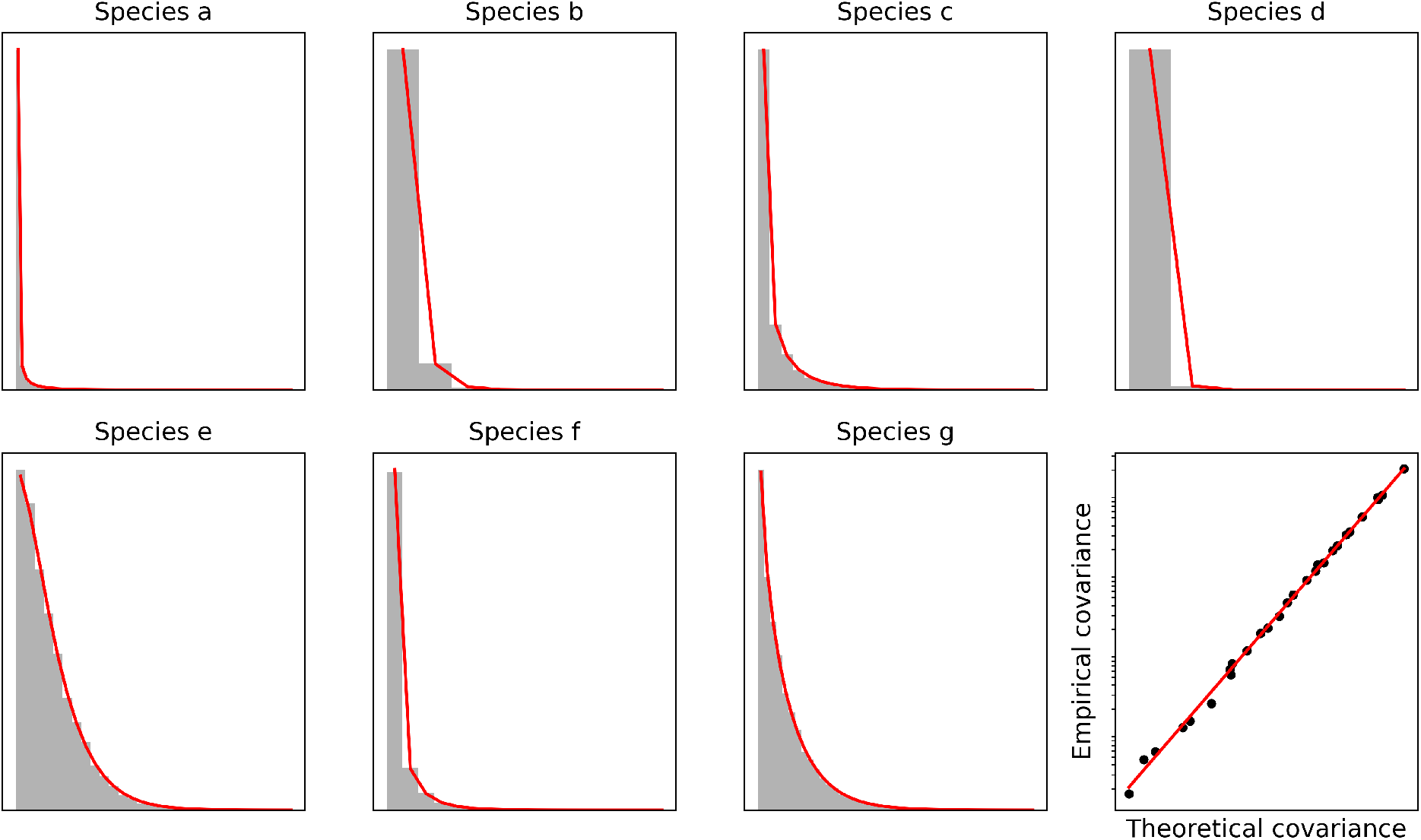
Simulation of the randomly generated graph model. Histograms: empirical results (10,000 simulations). Black dots: theoretical and empirical covariance. Red lines: theory.

### 2.6 Multi-gene systems

Although the CME solution is a general framework, the relevance to modern transcriptomic experimental data is tempered somewhat by the simplicity of the model. The model can describe the splicing cascade of a single gene, but does not naturally extend to multi-gene networks. Yet we know that genes often belong to co-expression modules that are identifiable by similarity metrics [3, 27]. Therefore, we are faced with the challenge of integrating multiple genes in a physically meaningful way.

Instead of building intractable “top-down” models that encode complex networks, we may build “bottom-up” models that extend analytical solutions. For example, we can consider sets of *synchronized* genes that experience bursting events at the same time. This model represents the bursty limit of multiple genes with transcription rates governed by a single telegraph process, up to scaling; a conceptually similar model has previously been used to describe correlations between multiple copies of one gene [28]. This model retains the appeal of physical interpretability – for example, gene modules may be regulated by the same molecule – but does not excessively complicate the mathematics, and offers an incremental step toward more detailed descriptions.

#### 2.6.1 Two-gene bursty model, no splicing

We begin by considering the instructive model of two genes influenced by the same regulator, with no downstream splicing. The burst processes are synchronized, but the burst sizes are not, and may indeed come from different distributions.

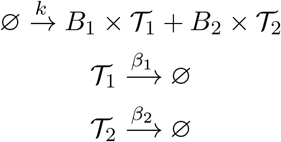

The following CME holds:

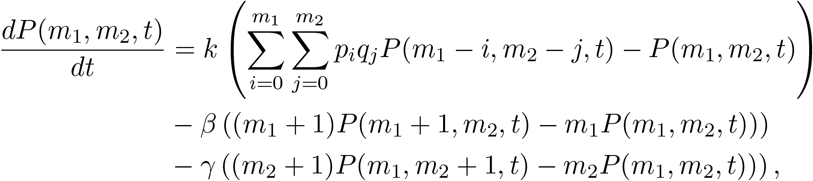

where *p*_*i*_ and *q*_*j*_ give the PMF weights of the burst size distributions that govern *B*_1_ and *B*_2_. We define the joint PGF

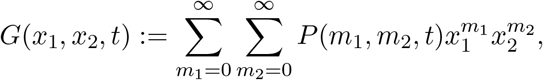

and recognize that the degradation terms have the familiar functional forms 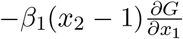 and 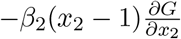. Therefore, considering the burst term:

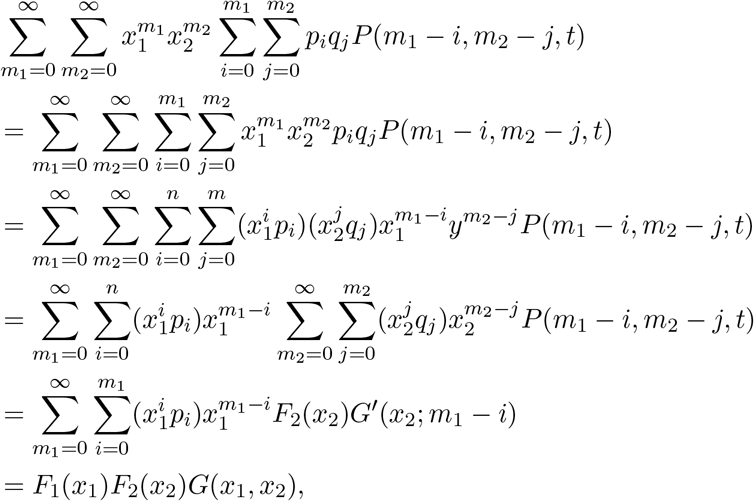

where *G′* is the conditional PGF assuming *m*_1_ − *i* molecules of 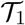 and *F*_*i*_ are the burst distribution PGFs; the final steps exploit the interpretation of the double sums as Cauchy products. This result implies:

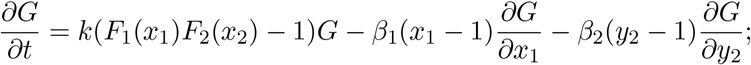

defining *G* = *e*^*ϕ*^, *u*_*i*_ = *x*_*i*_ − 1, and *M*_*i*_(*u*_*i*_) = *F*_*i*_(1 + *u*_*i*_):

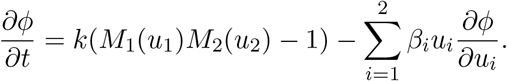

The PDE can be solved using the method of characteristics, yielding 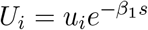:

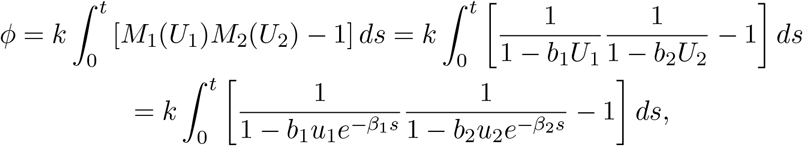

assuming, as before, that *B*_*i*_ ~ *Geom*(*b*_*i*_).

#### 2.6.2 *n*-gene bursty model with splicing

From the derivation above, four interesting properties stand out. Firstly, the functional form of *U*_*i*_ depends only on the details of the downstream processes; as shown, it can be easily computed for any DAG. Secondly, the derivation holds with no loss of generality for *any* number of genes, so long as the burst distributions are uncoupled, by repeated application of the Cauchy product formula.

Thirdly, even if they *are* coupled, the derivation still holds, but with intermediate conditional PGFs *F*_*i*_, ultimately yielding a joint PGF *F* that factorizes in the independent case. Finally, even though we have considered the case of disjoint DAGs, the results still hold if the gene products can ultimately converge. This becomes self-evident by identifying 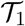 with 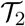 and setting *u*_1_ = *u*_2_, *b*_1_ = *b*_2_, and *β*_1_ = *β*_2_: the synchronized loci are merely multiple copies of the same gene, and produce bursts sampled from a *negative binomial* distribution, the sum of two identical geometric distributions. Therefore, in the most general case of arbitrary burst distributions and downstream splicing cascades, the factorial-cumulant generating function takes the following form:

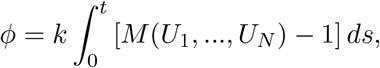

where *U*_1_, *..., U*_*N*_ are now the “*U*_1_” functions, *not necessarily distinct*, corresponding to the species produced by each bursty locus, and *M* is the joint MGF of the burst distributions. The single-gene results are recovered by marginalization.

#### 2.6.3 Examples of multi-gene systems

##### Uncorrelated gene loci

By the Cauchy-Schwartz inequality, all moments and cross-moments exist. The correlations are straightforward to compute. For example, we can consider the stationary covariance of gene products from two co-expressed genes, with disjoint downstream splicing processes. We marginalize over all other transcripts and consider the functional forms *Ul* = *ulψl* and *U*_*i*_ = *u*_*i*_*ψ*_*i*_, supposing that the respective average burst sizes are *b*_*l*_ and *b*_*i*_.

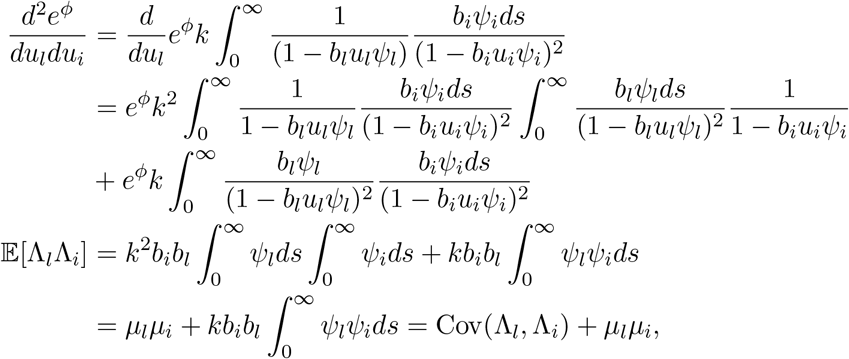

which implies that

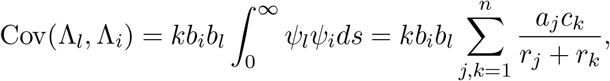

where *a*_*j*_ are the weights associated with *ψ*_*l*_ and *c*_*k*_ are the weights associated with *ψ*_*i*_, much as before. The covariance of the discrete process is identical. We reiterate that this expression *only* applies when the splicing graphs downstream of the gene loci are disjoint. The converse case requires slightly more unwieldy notation to account for, e.g., species *l* accessible from *N* source transcripts being associated with distinct coefficients 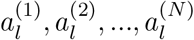, and does not appear to have a simple analytical expression totally agnostic of the number of loci and accessibility; nevertheless, it is easily tractable by appropriately defining the function sets 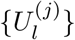 and 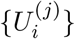 and using the procedure above.

With this solution in hand, we can revisit the two-gene problem with no splicing:

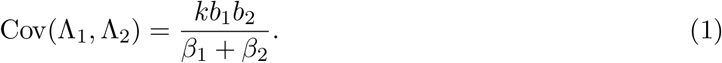

We are interested in standard summaries, such as the Pearson correlation coefficient:

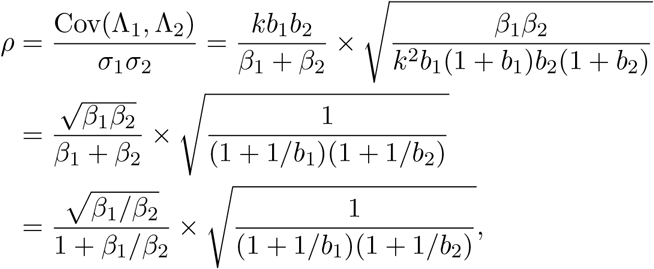

where we use the fact that the absolute timescale is immaterial. The first term achieves a global maximum of 1*/*2 at *β*_1_ = *β*_2_. The second is strictly smaller than 1, but asymptotically approaches 1 as *b*_1_, *b*_2_ jointly approach infinity. All downstream processes are stochastic and desynchronize molecular observables. Therefore, 1*/*2 is the supremum of gene-gene correlations in this class of models.

##### Fully correlated gene loci

Conversely, we can consider the two-gene problem assuming that the burst distributions are identical and fully correlated. Physically, this model may correspond to coupling of *initiation* processes, e.g. this may occur when two genes are controlled by a single promoter. This burst distribution has the following joint PGF:

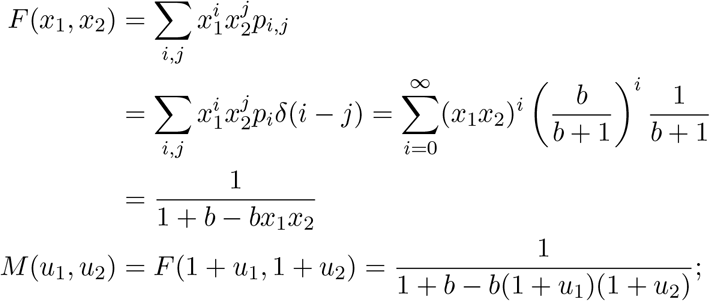

upon inserting the characteristics, we yield

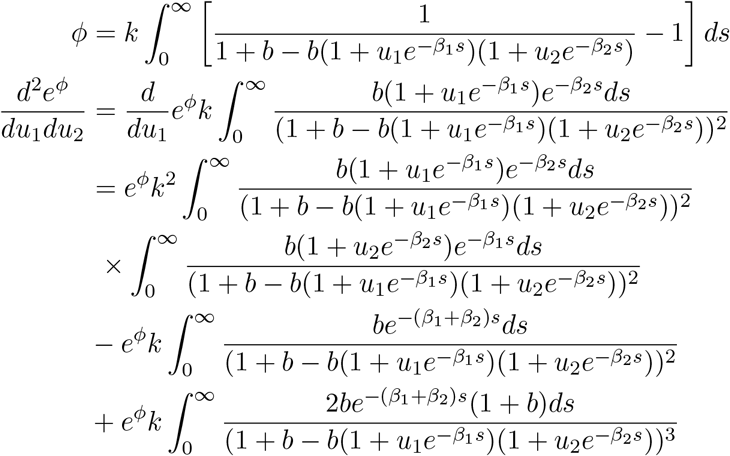

Plugging in *u*_1_ = *u*_2_ = 0,

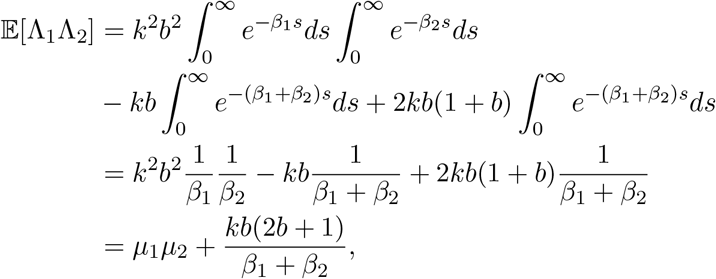

which yields the result

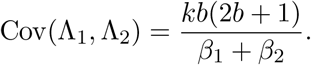

Therefore, the correlation is

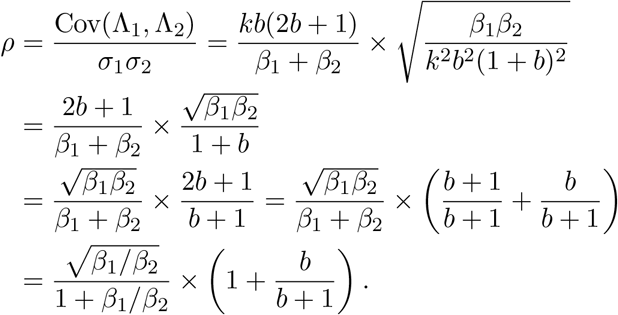

As in the case of uncoupled gene sizes reported in Equation 1, the first term is at most 1*/*2. The second term asymptotically approaches 2 as *b* → ∞. Therefore, there are no intrinsic model constraints on Pearson correlation coefficients of two gene products; constraints arise as the *effect* of the burst size correlation structure.

##### Anti-correlated gene loci

With these results in mind, we can consider the problem of describing genes with high *negative* correlations. As an illustration, we can consider two genes driven by a single telegraph process: gene 1 is on whenever gene 2 is off and vice versa; therefore, their respective products 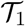 and 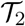 must have a negative correlation. However, only one of these genes has a well-defined bursty limit that is infinitesimally short on periods; the other will essentially be on all the time and transcribe constitutively, containing no mutual information about the burst timing. Nevertheless, putting aside the problem of positing a specific limiting mechanism, we can ask whether *any* joint burst distribution can produce negative correlations in molecule counts, *despite* perfect synchronization between burst events. Considering the cross moment of mRNA produced at two synchronized loci:

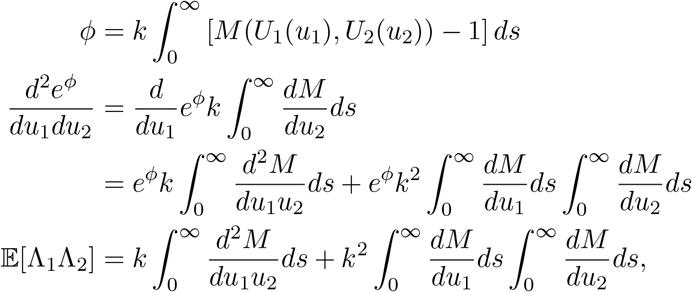

with the partial derivatives evaluated at *u*_1_ = *u*_2_ = 0. The second term matches *μ*_1_*μ*_2_, and is strictly positive. The first term is the integral of an exponentially discounted burst cross moment:

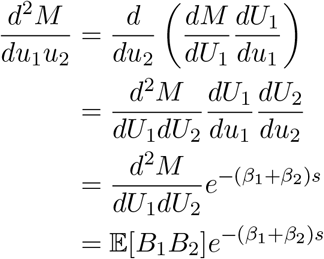

where the partial derivatives are yet again evaluated at *u*_1_ = *u*_2_ = *U*_1_ = *U*_2_ = 0, and *B*_1_ and *B*_2_ denote the SDE jump sizes at the two gene loci. By the definition of covariance:

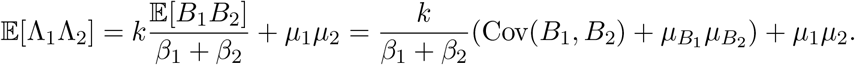

Now, supposing the correlation between the burst sizes is *ρ* ∈ [−1, 0), and considering the covariance between the transcripts:

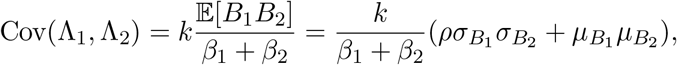

which achieves a minimum at *ρ* = −1. Thus, the covariance has a lower limit:

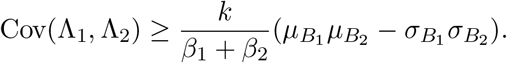

Without constructing the joint distribution explicitly, if we suppose the marginal discrete burst distributions are geometric – i.e., the jump sizes are exponential – then 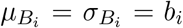, and the lower limit on covariance is zero. This means that negative correlations cannot possibly result from a model with geometrically distributed, synchronized jumps. However, other joint burst laws *can* produce negative correlations, as long as the population correlation coefficient is sufficiently negative and the burst distributions are sufficiently dispersed.

We can demonstrate the existence of processes with negative count correlations induced by synchronized burst events. First, we suppose that the marginal burst distributions are identical and described by a gamma law with shape *α* and scale *θ*, enforcing 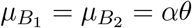 and 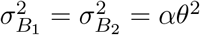. Therefore, the covariance of the Poisson intensities takes the following form:

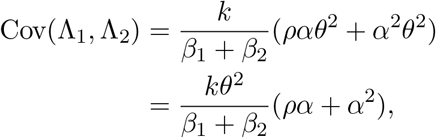

which achieves Cov(Λ_1_, Λ_2_) < 0 whenever *ρα* + *α*^2^ < 0. Therefore, for any *ρ* ∈ (−1, 0), every *α* ∈ (0, −*ρ*) meets this criterion.

It remains to confirm that a bivariate gamma distribution with a negative correlation can exist. Such a distribution was constructed by Moran, and permits all *ρ* ∈ (−1, 1) [29, 30]. Furthermore, a simple application of the Cauchy-Schwartz inequality yields [31]:

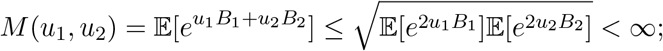

therefore, the joint MGF of the correlated bivariate gamma distribution is guaranteed to exist. This demonstrates the existence of continuous moving average processes with negative stationary correlation, driven by one Poisson process arrival process. Finally, the corresponding Poisson mixture has identical covariance, and must also have a negative correlation. Therefore, a CME with marginal negative binomial burst distributions and a carefully chosen joint structure can achieve negative molecular correlations, even if the bursts are synchronized.

##### Multi-gene dynamics emerging from fast processing

Interestingly, there is a set of single-gene systems that recapitulate the multi-gene functional form in the limit of fast splicing. Consider a source species 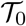, which is produced in geometrically distributed bursts and converted to species 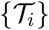, with *i* = 1, *..., N*, at rates *β*_*i*_. These transcripts are degraded at rates *γ*_*i*_.

Furthermore, suppose all of the 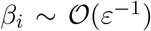 for small *ε*, i.e., the source transcript is extremely unstable. In this limit, 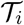 are produced with bursts of size *Bβ*_*i*_/*r*, where *B* is the underlying 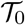 burst size, *r* ≔ Σ_*i*_ *β*_*i*_, and the ratio *β*_*i*_/*r* is *O*(1). We define corresponding *weights w*_*i*_ ≔ *β*_*i*_/*r*; by definition, Σ_*i*_*w*_*i*_ = 1. This yields:

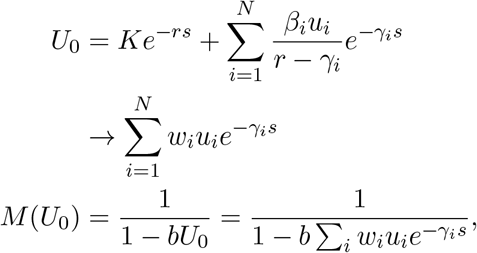

where *M* is recognizable as the joint MGF of a perfectly correlated *N*-variate exponential distribution with marginal distributions *B*_*i*_ = *w*_*i*_*B*:

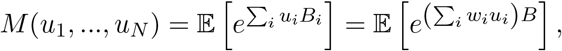

i.e., the univariate exponential MGF evaluated at 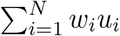.

The corresponding discrete PGF is:

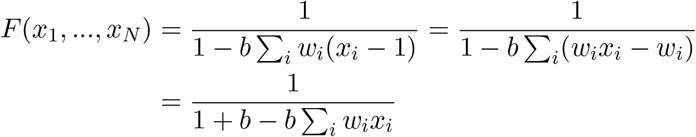

As seen above, this is *not* the perfectly correlated multivariate geometric distribution: the stochasticity of the reaction channel selection is non-negligible. Instead, we can construct a distribution over {0, 1}^*N*^, with the probability of state *δ*_*ij*_ (i.e., the vector contains a one at position *i*) set to *w*_*i*_. It is easy to see that the generating function of the random variable *Z* ∈ {0, 1}^*N*^ takes the form derived above:

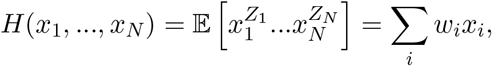

which amounts to *F*(*x*_1_, *..., x*_*N*_) = *F*(*H*(*x*_1_, *..., x*_*N*_)); i.e., the effective gene dynamics are described by a compound distribution.

Upon inserting the characteristics and selecting a set of two genes (arbitrarily indexed by 1 and 2), we yield:

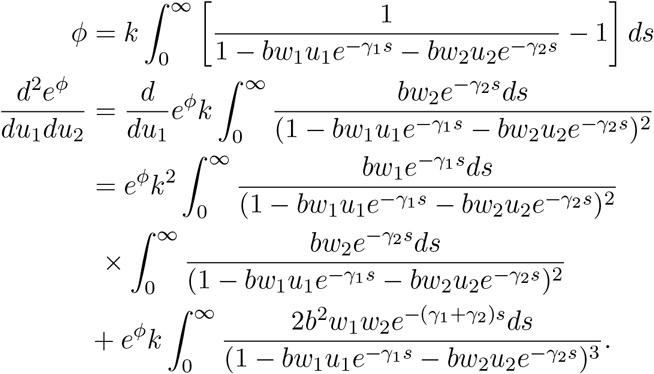

Plugging in *u*_1_ = *u*_2_ = 0,

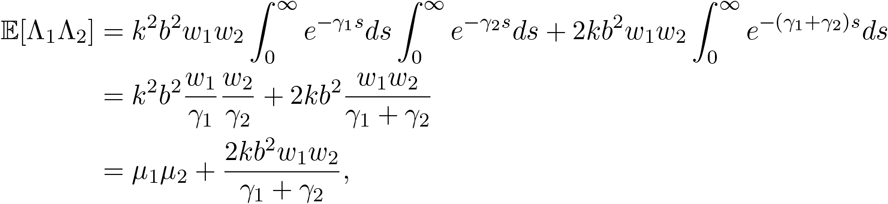

since marginalization with respect to any gene recovers the univariate geometric burst distribution with scale *bw*_*i*_, which immediately yields *μ*_*i*_ = *kbw*_*i*_/*γ*_*i*_. This implies:

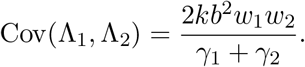

From the marginal results, we still have 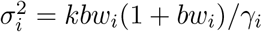. This yields:

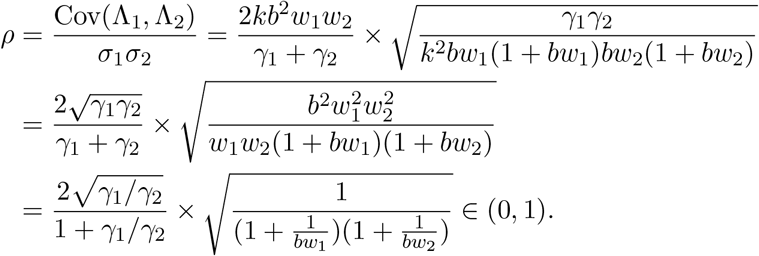

Therefore, fast processing of the source transcript yields dynamics equivalent to a class of multi-gene models, with positive, but otherwise unconstrained, correlation between the downstream species.

### 2.7 Transient gene dynamics

Thus far, we have primarily focused on stationary systems with time-independent parameters. Nevertheless, there are classes of physiological phenomena, such as differentiation and cell cycling, where transient behaviors are crucial, particularly since these processes occur on timescales comparable to the mRNA lifetimes [1, 32]. Usually, the regulatory events underpinning these processes are modeled by variation in DNA-localized transcriptional parameters [33–36].

By examination of the generating function relations, it is easy to see that the current framework is trivial to extend to *any* deterministic variation in *k* and *M*:

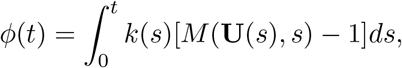

where we have adopted the shorthand **U** for the set of exponential sums *U*_1_(*s*), *..., U*_*N*_(*s*) governing each burst product of the *N* loci. Therefore, burst frequency, burst size, and even the number of synchronized gene loci per cell can vary, continuously or discontinuously.

If the reaction rates within **U** change over time, the generating function PDE then becomes intractable for all but the simplest models, such as piecewise constant. The gene locus parameters *b* and *k* can also vary stochastically. In principle, certain models can be solved by defining an appropriate SDE-CME system. However, such dynamics are generally challenging to treat without recourse to numerical ODE solvers. Further, we restrict our analysis to systems tractable *via* the PGF, which cannot be applied to continuous-valued parameters.

## 3 Applications to sequencing data

The current framework provides quadrature-based solutions to the forward problem of PMF prediction for a broad set of transcriptional processes. More broadly, we would like to treat the *inverse* problem of identifying parameters from sequencing data. A wide range of statistical approaches are available; however, in practice, even the simplest, ergodic version of the the inverse problem depends on the following prerequisites:

1. Single-cell, single-molecule data for a set of cells in local equilibrium. This information permits the application of the ergodic model.
2. Full annotation of intermediate transcripts, *including causal relationships*, such as the splicing graph and the identities of degraded molecules. This information permits mapping between experimental data and theoretical quantities.
3. Transcriptome-wide quantification of all transcripts, ideally unbiased and fully saturated – or, at the very least, imperfect quantification combined with a statistical model of sequencing. This information permits the inclusion of technical noise.

No perfect experimental solution exists. The collection of single-cell, single-molecule data is enabled by barcoding [37]. Characterization of splicing graphs has been treated in experimental [38, 39] and computational [40] contexts. However, these necessarily rely on comprehensive single-molecule annotations – which distinct intron/exon combinations occur? – which have only become feasible on a genome-wide scale since the introduction of molecular barcoding. Fully saturated sequencing is infeasible due to cost, and potentially due to thermodynamic constraints. Finally, standard sequencing capture protocols are, by design, biased toward polyadenylated regions [37]. This effect has been exploited to capture natural intronic sequences [1] and synthetic antibody-conjugated oligonucleotides [41, 42], but the quantitative effects of these biases are as of yet unclear. Naturally, these data quality challenges are compounded with standard statistical challenges, such as the often considerable computational expense of determining confidence regions.

In spite of these challenges, we *can* immediately apply some of the simpler theoretical results to transcriptomic datasets.

We used long-read sequencing data generated by FLT-seq (full-length transcript sequencing by sampling) [43]. In brief, the method synthesizes a cDNA library using 10X gel beads and primers, amplifies it, then applies nanopore sequencing to generate long reads with associated cell and molecule barcodes. We obtained a mouse stem cell dataset (498 barcodes), filtered for the activated subset (136 barcodes), and identified the 1000 highest-expressed genes.

We used *gffutils* 0.10.1 to construct a database of identified intermediate and terminal isoforms, based on the accompanying annotations generated by the computational pipeline *FLAMES*(full-length analysis of mutations and splicing). We used the *intervaltree* 2.1.0 Python package to split the transcript-specific exons into elementary intervals, then used graph tools from the *NetworkX* 2.5.1 Python package to identify “root” transcripts that cannot be obtained by removing regions of any other transcript.

The presumed source transcript covering the entire locus was not observed for any of the genes. As their exonic patterns are mutually exclusive, we model each gene’s root transcripts as products of a single rapidly processed source species, as in Section 2.6.3. The theoretical results immediately imply that the root transcripts must be distributed according to the negative binomial distribution.

Therefore, by fitting the transcript distributions, we can estimate effective burst sizes *bw*_*i*_ and unitless efflux rates *γ*_*i*_/*k*. These marginal parameters can be plugged into the formula for Pearson correlation derived in Section 2.6.3, and compared to the empirical correlation coefficients. The theoretical correlations are to be understood as upper limits, as unobserved intermediate processing steps and technical noise inevitably reduce transcript–transcript correlations.

We fit the root transcript marginal distributions using the *statsmodels* 0.10.2 Python package. One gene was rejected outright due to failure to construct elementary intervals. 732 transcripts were rejected due to underdispersion (mean higher than variance), as they do not possess valid maximum likelihood estimates. 1290 transcripts were rejected due to poor fit (failure to converge or excessively sparse data, with all but one rejected transcripts having average expression below 1 mRNA per cell). This analysis produced a set of 4362 transcripts and 12480 nontrivial correlation matrix entries.

Theoretical and observed correlation coefficients are shown in Figure 6. 11894 of the theoretical correlations (95.3%) are higher than the observed ones, whereas 586 (4.7%) are lower. Furthermore, the theoretical correlations are clearly nontrivial; all are less than unity. The results suggest that the model is not sufficient to recapitulate the full dynamics, but *does* provide an effective theoretical constraint. We anticipate that the lower observed correlations emerge from a contribution of technical noise in the sequencing process, suboptimal inference from the marginals, correlation-degrading stochastic intermediates, and model misidentification. The final effect is particularly plausible for the cluster evident on the left side of the figure, as the fast-processing model cannot account for negative transcript–transcript correlations.

**Figure 6:**
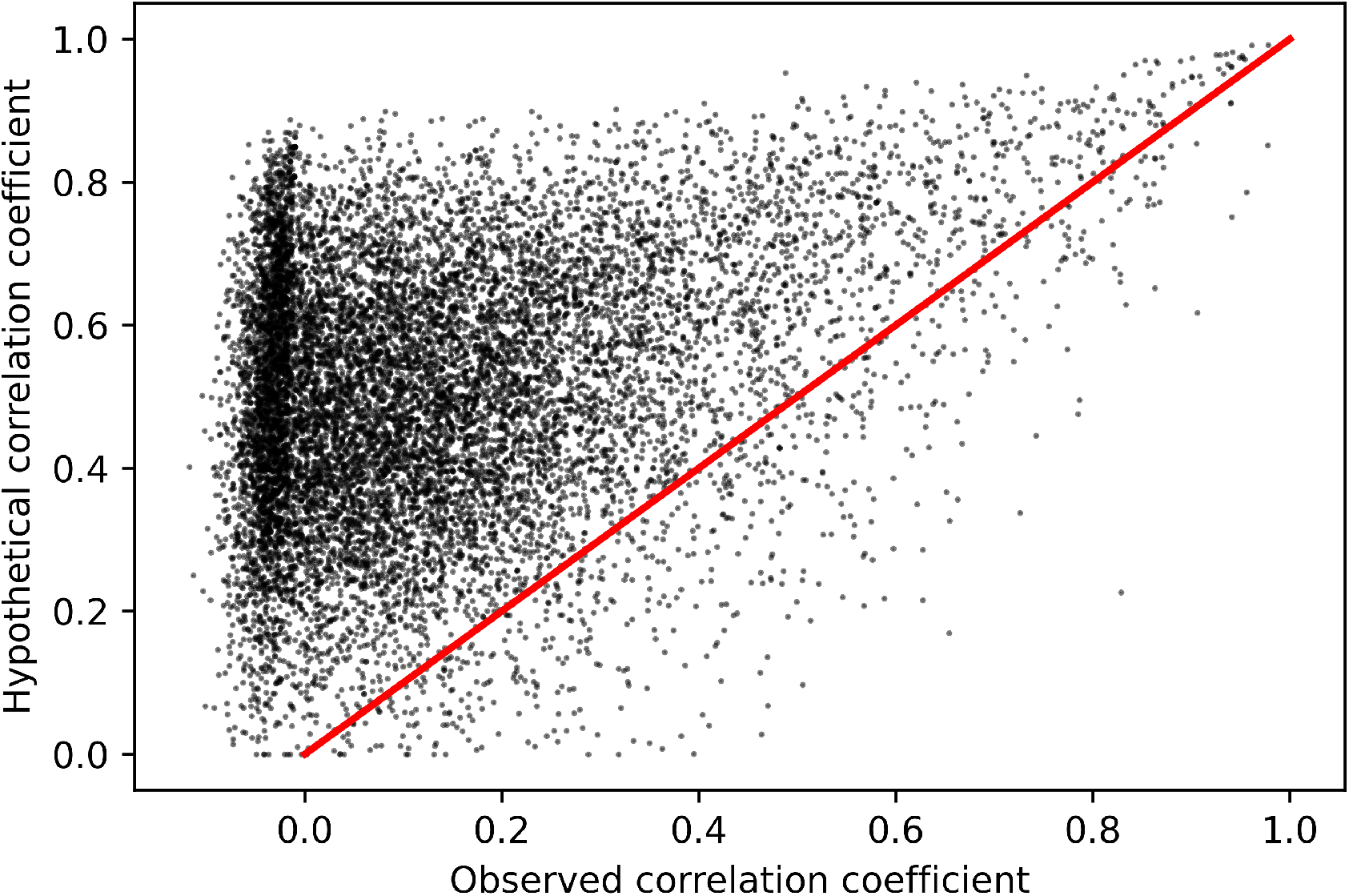
Theoretical Pearson correlation coefficients inferred from marginal distributions, compared to empirical correlations. The theoretical model generally overestimates transcript–transcript correlations, and rarely underestimates them, which is consistent with the interpretation of the theoretical correlation as a nontrivial upper limit.

## 4 Results and discussion

We have described a broad extension of previous work pertaining to monomolecular reaction networks coupled to a bursty transcriptional process. In particular, by exploiting the standard properties of reaction rate equations, we have demonstrated the existence of all moments and cross-moments. Further, we have derived the analytical expressions for the generating functions and demonstrated their existence. The following expression gives the general solution for the factorial-cumulant generating function:

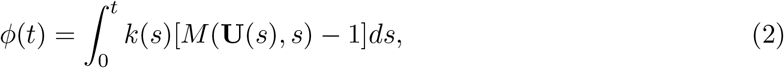

where **U** is a set of functions associated with the gene products, computable by the procedure in Section 2.2.3, and *M* is a joint generating function governing the bursting dynamics.

Under the mild assumptions of finite-activity bursting and Markovian downstream processes, the expressions hold for arbitrary directed acyclic graphs of splicing and degradation, coupled to an arbitrary number of gene loci with arbitrary, and potentially correlated and time-dependent, burst dynamics. In fact, we explicitly consider the problem of modeling multiple synchronized genes and find that geometric burst size coordination is *required* to achieve transcript count correlations *ρ >* 1/2. Furthermore, we test whether negative correlations are feasible under the assumption of synchronized bursts at multiple gene loci, and find that *ρ <* 0 are impossible with geometric bursts, but *can* be achieved with negative binomial bursts. These results substantially constrain and inform the space of models that can recapitulate the combination of bursty dynamics [11] and high absolute gene-gene correlations [3, 27] observed in living cells.

We compared the theoretical constraints with experimental data generated by FLT-seq, a recent long-read, single-cell, single-molecule sequencing method. Investigating a set of 4362 transcripts, we found that the constraints were met for 95.3% of the transcript–transcript correlations. Nevertheless, the model was insufficient to recapitulate the precise quantitative details, suggesting that more detailed biophysical models of splicing networks and technical noise are necessary.

Although we primarily focus on distributions, with an eye to inference from atemporal data, the solution is robust in several complementary dimensions. Firstly, the upper limit of the integral in Equation 2 is arbitrary, and can be evaluated up to a finite time horizon to yield a transient distribution. Secondly, the formulation of the problem presupposes *m*_*i*_(*t* = 0) = 0 for all species *i*. However, if nonzero molecule counts are present at *t* = 0, it is straightforward to compute the resulting log-PGF via 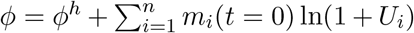, where *ϕ*^*h*^ is the result for *m*_*i*_(*t* = 0) = 0 and the *U*_*i*_ are species-specific exponential sums. This relation produces arbitrary *conditional* distributions, as derived elsewhere [44]. Therefore, the current approach can be used to compute the likelihoods of entire trajectories of observations, and thus perform parameter estimation on live-cell data.

The computational complexity of this procedure is 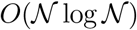 in state space size. However, this complexity ostensibly arises from the *n*-dimensional inverse Fourier transform. We expect the time complexity to be 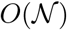 in the practical regime, and irrelevant outside due to the *space* complexity of holding an array of size 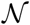 in memory for the inverse Fourier transform.

Curiously, this class of analytical solutions to reaction networks can be adapted to a subset of diffusion problems. General diffusion on a multidimensional lattice is not directly solvable, because it violates the assumption of acyclic graph structure. However, percolation through a directed acyclic graph coupled to a source and a set of sinks can be described using the current mathematical formalism. Further, such a percolation can represent the incremental movement of RNA polymerase along a DNA strand, integrating discrete copy number statistics with submolecular details in a single analytical framework [45, 46].

We do not treat the aforementioned auxiliary degenerate solutions that arise when *β*_*i*_ = *β*_*j*_ for some *i, j*. However, Section 2.4 guarantees the existence of these solutions and all moments: the procedure does not rely on a particular functional form of *ψ*_*i*_, only its existence, which is implied by the test subordinator *F*(*x*) = *x* matching the constitutive case [14].

Finally, we briefly touch upon the class of delay chemical master equations, and survey several recent advances in the field in Section S1.1. Due to the non-Markovian nature of delayed systems, general probabilistic solutions are rare [47] and represent an open area of study. In our discussion, we motivate delays as a limit of numerous, fast isomerization processes, and clarify the challenges inherent in applying the analysis of delay CMEs to bursty systems.

## 5 Appendix: ODE solution

The solution to the generic ODE 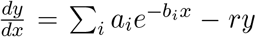 is 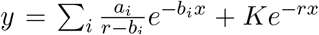. Using the initial condition *y*(0) = *ξ* yields 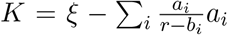. Consider an efflux rate *r* = Σ_*j*_*β*_*j*_+*γ*, i.e., a set of isomerization pathways and a single degradation pathway from the species in question. Care must be taken when the downstream paths converge (as in the case of the two-intron system): duplicate terms in product characteristics *U*_*j*_ need to be aggregated, with Σ_*j*_*β*_*j*_ *U*_*j*_ rewritten as 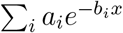. This is computationally straightforward to do by choosing appropriate data structures, as with the matrix *A* in the implementation of the DAG algorithm.

## 6 Code availability

Python notebooks that can be used to reproduce Figures 5 and 6 are available at https://github.com/pachterlab/GP_2021_2.

## 7 Acknowledgments

The DNA illustration used in Figures 1 and 4, modified from [44], is a derivative of the DNA Twemoji by Twitter, Inc., used under CC-BY 4.0. The directed acyclic graph generation code was adapted from the IPython Parallel reference documentation: https://ipyparallel.readthedocs.io/en/latest/dag_dependencies.html. The yellow and gray colors used in Figures 1, 3, and 4 are the Pantone Colors of the Year 2021, PANTONE 17-5104 Ultimate Gray and PANTONE 13-0647 Illuminating. G.G. and L.P. were partially funded by NIH U19MH114830.

## S1 Supplementary Note

### S1.1 Delay chemical master equations

In the current supplement, we detour from the Markovian framework to consider *delay* systems, which have deterministic, rather than stochastic, state transitions. Certain degenerate cases – for example, the problem of incremental, linear movement with identical transition rates – directly bear upon the class of delay chemical master equations (DCMEs). As an example, we can model the simple linear chain of reactions with constitutive production

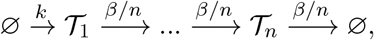

the total delay between production of 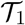 and degradation of 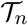 is Erlang-distributed, with shape *n* and rate *β*. As *n* → ∞, the Erlang distribution reduces to a point mass at *β*^−1^ ≔ *τ*. This implies that we can treat an *aggregated* species 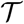, produced at rate *k* and degraded after a deterministic delay *τ*:

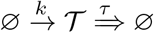

This is precisely the “linear chain trick” introduced by MacDonald in 1978 [48]. The study of delayed dynamical systems, such as delay differential equations, dates back to the eighteenth century [49], with cornerstone biological models by Lotka and Volterra [48, 49]. Recent work has focused on developing exact solutions [47, 50, 51] and simulation methods [52, 53]. In particular, studies by Lafuerza and Toral [54, 55] report full analytical solutions for constitutive systems with isomerization, while a contemporary study by Jia and Kulkarni [56] reports lower moments for a system with bursty mRNA production and catalysis.

Unfortunately, applying these methods to bursty systems is challenging, and all but the simplest problems are intractable. As an illustration, we consider the constitutive two-stage system described by by Lafuerza and Toral [54], and discuss the challenges of extending it to include bursty production. If we assume that no stochastic degradation reactions occur, the reaction equations and generating function relations take the following form:

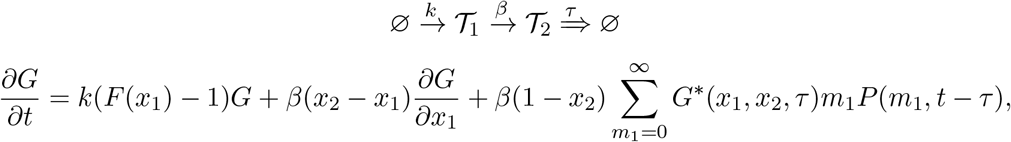

where *G*\* is a conditional generating function for an auxiliary *non-degrading* system, initialized at *m*_1_ − 1 molecules of the parent transcript 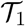. This auxiliary system has no degradation reactions, and allows us to incorporate the non-Markovian effects of delays. Assuming constitutive production, and using the shifted variables *u*_*i*_ for convenience, we find:

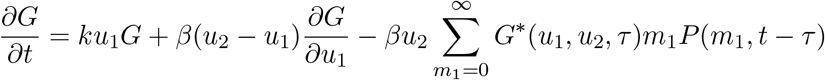

The final term is *not* proportional to *G*, so no convenient exponential *ansatz* is available. However, the sum affords an alternative representation, which exploits the separability of the initial condition and the dynamics on [0, *τ*]:

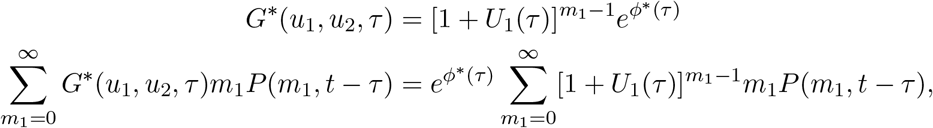

where *ϕ*\* is the factorial-cumulant generating function of the auxiliary system, started at zero molecules. This sum may be treated as the first derivative of the stationary 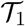 PGF, evaluated at 1 + *U*_1_, where *U*_1_ is a function computed by solving the non-degrading system with the method of characteristics.

We start by computing the auxiliary *U*_1_ by using the method of characteristics and enforcing *U*_2_(0) = *u*_2_ and *U*_1_(0) = *u*_1_.

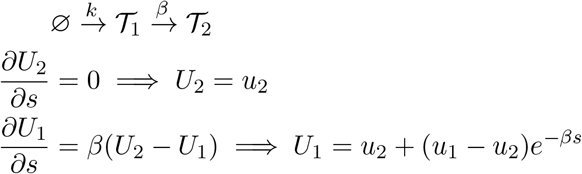

Now, we compute the generating function of the subsystem:

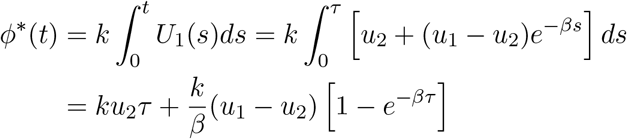

We compute the derivative of the 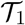 Poisson PGF:

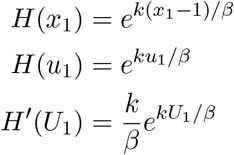

This construction is slightly simpler than in the original: we do not use the full time-dependent Poisson distribution, but presuppose that the system starts with 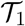 in equilibrium. Since it approaches this distribution exponentially fast regardless of initial conditions, the error is minimal, and the simplification eliminates the time dependence in the degradation term.

Plugging in and evaluating the non-Markovian term:

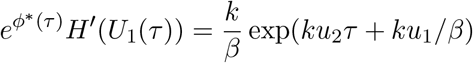

Finally, considering the full generating function expression:

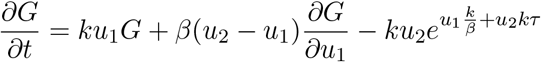

Lafuerza and Toral report a solution [54], though computed through an *ansatz* rather than directly – this PDE is not quite as simple as that of the Markovian system. We restrict ourselves to the stationary solution, which can be solved with a rather mechanical application of the integrating factor method, or by noticing that the uncorrelated Poisson PMF solves the equation:

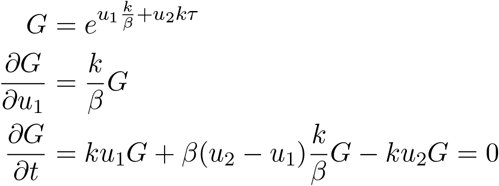

Of course, this result can be derived just as well without writing down anything at all – by using the standard results for constitutive production [14], the linear chain trick, and the fact that sums of Poisson random variables are Poisson. However, the rigorous approach can be used to treat more general systems. In particular, we attempt to solve the delayed analog of the two-stage bursty system [10]:

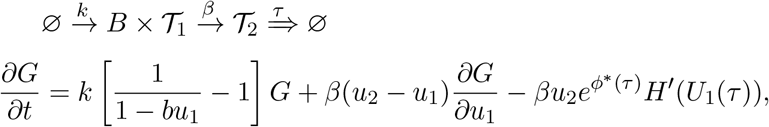

where the auxiliary system is now bursty.

First, we compute the factors of the non-Markovian term. The PGF derivative is found by evaluating the 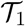 marginal:

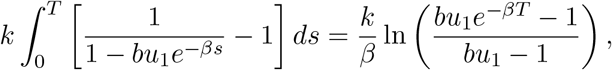

which coincides with the relevant result for the gamma Ornstein–Uhlenbeck SDE [21]. However, this form is needlessly challenging to work with, and it is more straightforward to assume *T* ≫ 0, or the system starts in the equilibrium distribution of 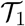. Again, due to exponential convergence, the error is minimal. Differentiating with respect to *x*_1_ = *u*_1_ + 1:

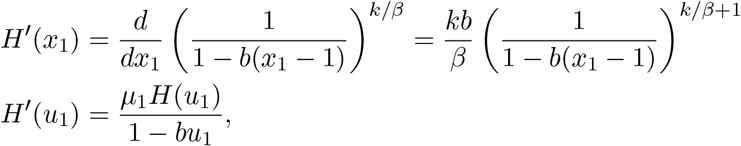

where we define *μ*_1_ ≔ *kb/β* for simplicity. This yields a straightforward expression for the summation:

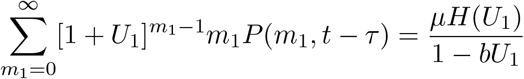

We reuse *U*_1_ and *U*_2_ from the derivation of the constitutive system, as the downstream components of the auxiliary systems match:

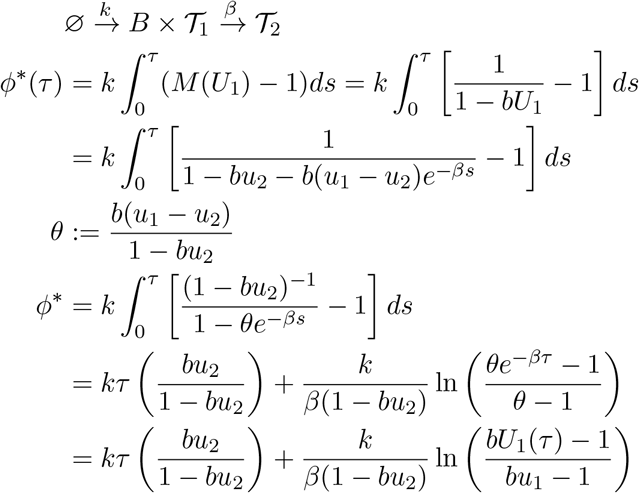

which follows from the derivation of the PGF of the nascent marginal.

Now, considering the full generating function relation:

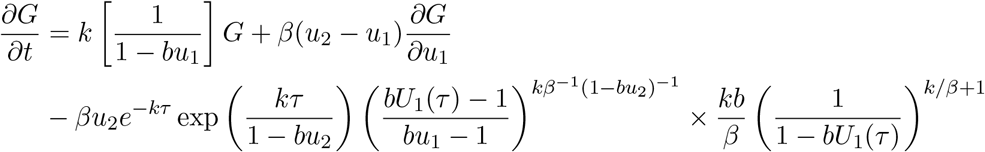

This PDE is not easily tractable by standard analytical or numerical methods. The form of the equation is rather complicated and not amenable to analysis by characteristics. In principle, a numerical PDE or ODE solver can be used: we may fix *u*_2_ and solve for *G*(*u*_1_, *u*_2_) over a mesh of *u*_1_. By repeating this for many values of *u*_2_, we can compute the Fourier transform of the joint distribution. However, this requires solvers that can integrate over the complex plane, as well as initial conditions *G*(0, *u*_2_) for each *u*_2_. These are the very values we seek, so even numerical approaches require some ingenuity.

In short, the stochastically delayed systems reduce to deterministically delayed systems in some well-studied regimes. However, in spite of the formal connection between the CME and the DCME, the former is far simpler to analyze: the DCME is non-Markovian, and generally resistant to exact analysis. Although much recent progress has been made, regulated transcriptional systems do not yet have full probabilistic solutions.

